# Single Cell Portal: an interactive home for single-cell genomics data

**DOI:** 10.1101/2023.07.13.548886

**Authors:** Leyla Tarhan, Jon Bistline, Jean Chang, Bryan Galloway, Emily Hanna, Eric Weitz

## Abstract

Single-cell omics research has the power to leave a deep impact on modern healthcare. Sharing data widely and freely advances this progress in both the academic and clinical spheres. We developed the Single Cell Portal (SCP) to maximize the impact of this work. SCP enables data sharing, supports dynamic results visualization, and facilitates scientific exploration across a large repository of single-cell datasets. SCP’s data contributors maintain full control over how their data are shared and presented, without requiring web development expertise. Finally, SCP supports the entire lifecycle of a research project, from sparking an idea, to fine-tuning the data with collaborators, to sharing results in an accessible and interactive way. This paper highlights the most valuable ways in which SCP helps to advance single-cell research.

## INTRODUCTION

Since its development ([Tang et al. 2009](https://www.nature.com/articles/nmeth.1315)), single-cell omics research has seen an exponential growth and has achieved far-reaching scientific effects (**Figure 1**). Recent research from this field has taught us about the mechanisms behind many systems and diseases, from T-cells in COVID-19 to the structure of the healthy mouse brain (e.g., [Kolodziejcyk, 2015](https://www.cell.com/cell-stem-cell/fulltext/S1934-5909(15)00418-X?_returnURL=) Kolodziejcyk, 2015 [Suvà & Tirosh, 2019](https://linkinghub.elsevier.com/retrieve/pii/S1097276519303569); [Ziegler et al., 2020](https://www.cell.com/cell/fulltext/S0092-8674(20)30500-6?_returnURL=)Ziegler et al., 2020; [Kozareva et al., 2021](https://www.nature.com/articles/s41586-021-03220-z)). This work also has a large social impact, as understanding the basic mechanisms behind diseases helps us develop better clinical treatments (e.g., [Majumder et al., 2022](https://onlinelibrary.wiley.com/doi/10.1002/jmv.27558)).

**Figure 1.**
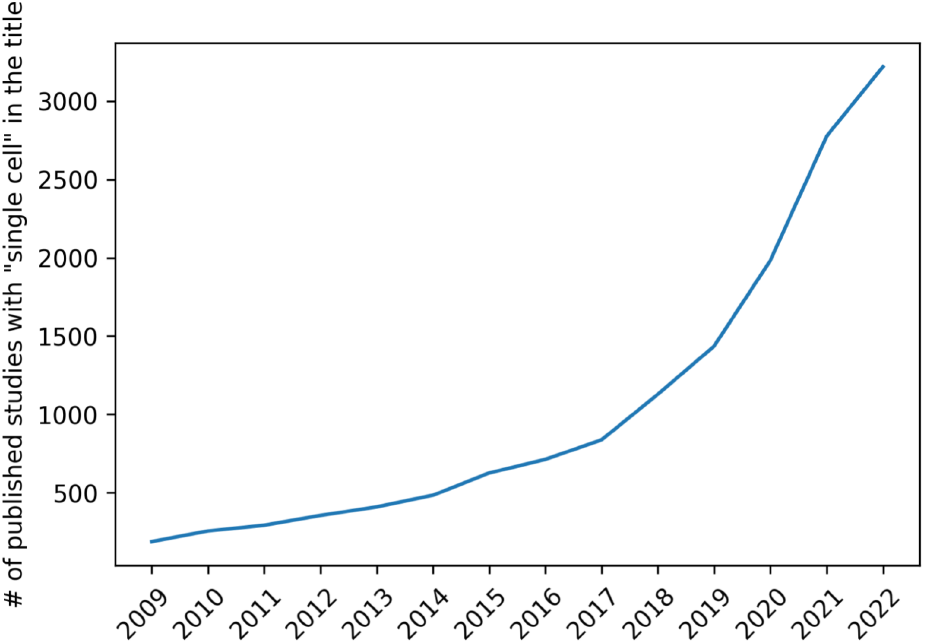
The number of papers with “single cell” in the title has increased exponentially since the method was first described in [Tang et al., 2009](https://www.nature.com/articles/nmeth.1315). Data from PubMed.

These advances in single-cell science are fueled by technological advances that allow researchers to retrieve and understand detailed sequencing information from large-scale data ([Blainey & Quake, 2014](https://www.nature.com/articles/nmeth.2783)). Several groups have developed platforms to host, share, and explore this data to accelerate single-cell science’s progress. These include [CellxGene](https://cellxgene.cziscience.com/) (Li et al., 2020; [Legill et al., 2021](https://www.biorxiv.org/content/biorxiv/early/2020/08/31/2020.08.28.270652.full.pdf)), [gEAR](https://umgear.org/) ([Orvis et al., 2021](https://www.nature.com/articles/s41592-021-01200-9)) and the associated [NeMO Analytics portal](https://nemoanalytics.org/) ([Ament et al., 2023](https://doi.org/10.1093/nar/gkac962)), the [Single-Cell Expression Atlas](https://www.ebi.ac.uk/gxa/sc/home) ([Papatheodorou et al., 2019](https://academic.oup.com/nar/article/48/D1/D77/5609521)), the [GTEx portal](https://gtexportal.org/home/) ([Carithers & Moore, 2015](https://www.ncbi.nlm.nih.gov/pmc/articles/PMC4692118/); [Carithers et al., 2015](https://www.ncbi.nlm.nih.gov/pmc/articles/PMC4692118/)), and [HuBMAP](https://hubmapconsortium.org/) ([Snyder et al., 2019](https://www.nature.com/articles/s41586-019-1629-x)).

Among the diverse features that differentiate these single-cell data platforms, the most important is the degree to which data contributors can customize their data’s presentation, while maintaining the data’s findability and reusability. On one hand, dataset-specific platforms developed with tools such as [ShinyCell](https://academic.oup.com/bioinformatics/article-abstract/37/19/3374/6198103) ([Ouyang et al., 2021](https://academic.oup.com/bioinformatics/article/37/19/3374/6198103)) offer interactive visualizations that allow researchers to control exactly how their data are presented. These visualizations make the data accessible to both computational and non-computational researchers. However, usage of these platforms tends to be restricted to fairly domain-specific niches of the single-cell research community. Under this model, it is also difficult for interested researchers to discover related datasets. The challenges of scaling these platforms for larger datasets also make this approach less adaptable as the field moves toward larger, atlas-style datasets. On the other hand, data repositories such as NIH’s Gene Expression Omnibus ([Edgar et al., 2002](https://academic.oup.com/nar/article/30/1/207/1332640)) allow researchers to share their data with relatively little effort, but offer very limited, if any, data visualization tools that could facilitate data exploration. As a result, researchers who are interested in a dataset have to download and analyze the data locally. As a result, this approach silos the data analysis process and puts up extra hurdles for researchers with less computational expertise.

Single Cell Portal (SCP) facilitates single-cell research by sitting in between these extremes. SCP (https://singlecell.broadinstitute.org/single_cell) is an open-access, web-based portal for sharing and exploring single-cell genomics data. It is self-service, meaning that any researcher can contribute data to the Portal. Once these data are uploaded, SCP renders interactive visualizations that allow other researchers to easily explore these data, even without a computational background. Data contributors can customize these visualizations without having to build them from scratch. In this way, SCP balances the tradeoffs between customizability, findability, and reusability. In addition, SCP is supported by a performant computational infrastructure, which allows visualizations to scale with growing datasets. It also collects a large library of open-access single-cell datasets, making it easy to search across many studies and then dive deeply into the most interesting ones.

These qualities allow SCP to support the full lifecycle of single-cell research, from the inception of an idea, to fine-tuning the data, to publishing and disseminating the results. In the following sections, we detail how engagement with SCP supports each of these research stages, and show how sharing data on SCP multiplies the impact of single-cell research studies.

## HOW SCP SUPPORTS SINGLE-CELL RESEARCH

SCP provides crucial support for every stage throughout the lifecycle of single-cell research (**Figure 2**). This engagement cycle begins when a researcher uses SCP’s interactive visualization tools to explore existing datasets, which may help them refine their scientific questions and experimental framework (**Figure 2a**). After the researcher has collected and cleaned a new dataset, they can deposit it on SCP in a private study that can be shared with collaborators to examine and refine the data (**Figure 2b**). During this refinement process, researchers can also compare patterns in the new dataset against SCP’s repository of public datasets (**Figure 2c**). When the data are ready to be shared with peer reviewers for a manuscript, SCP allows data contributors (“study owners”) to share their data with anonymous reviewers in a “view-only” mode (**Figure 2d**). When the data are ready to be released to the wider research community, the SCP study can be switched to ‘public’ mode (**Figure 2e**). Making the data available in an explorable form may increase the work’s impact. To further increase this impact, researchers can group their studies into a curated “study collection” (**Figure 2f**). Just as research begets more research, the cycle of engagement with SCP begins again as researchers explore more datasets, which may spark more ideas. The following sections describe these steps in the SCP engagement cycle in more detail, and show the impact of depositing data on the Portal.

**Figure 2:**
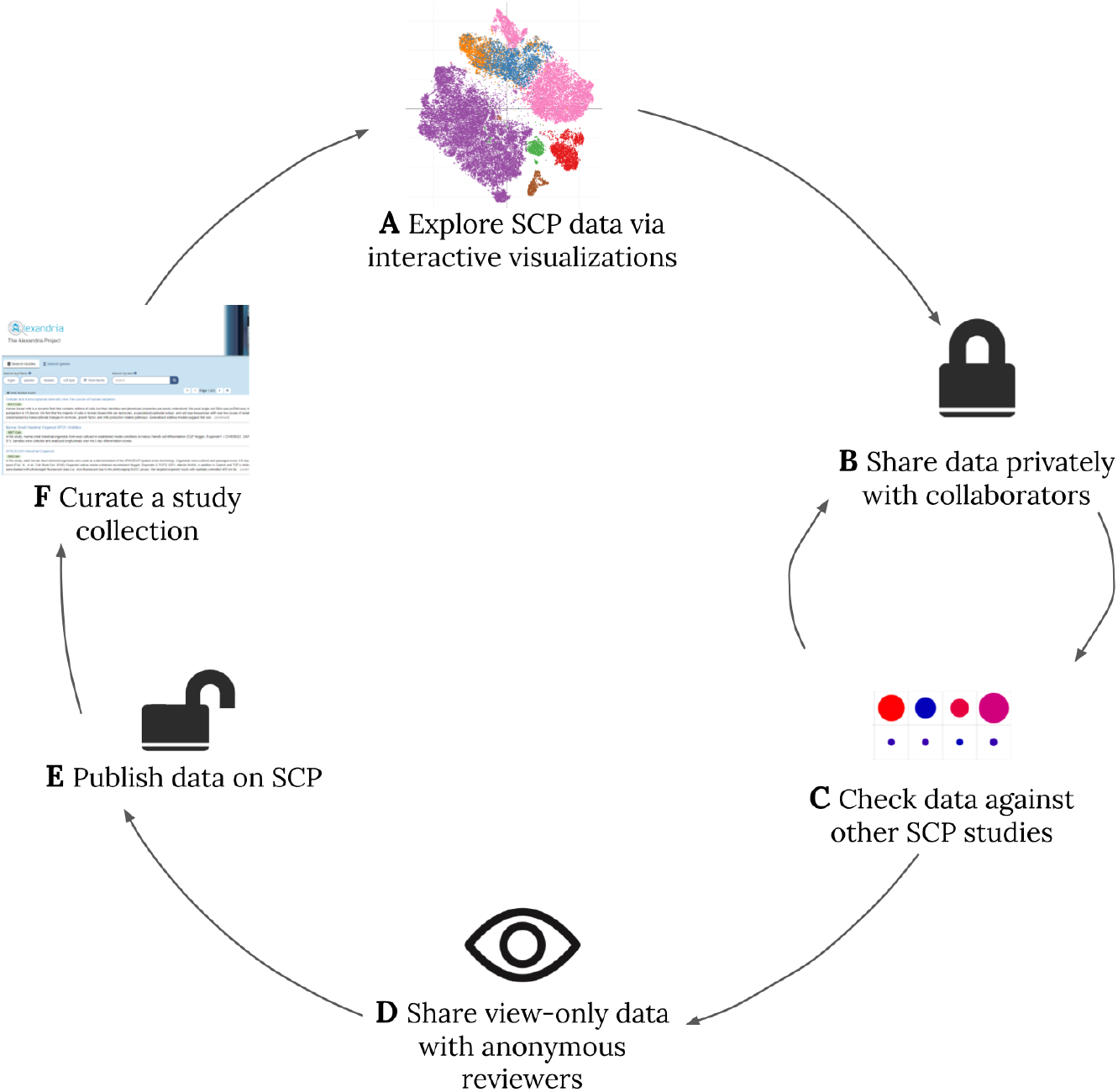
The Cycle of Engagement with Single Cell Portal. We outline a typical cycle of engagement with Single Cell Portal, which supports the larger cycle of single-cell research. (A) SCP provides an accessible way for researchers to explore other single-cell datasets, which may help refine hypotheses and ideas for experiments. (B) Once a new dataset has been collected and cleaned, it can be shared with collaborators as a private study. (C) To verify cell type annotations or other patterns in the data, researchers can check for similar patterns in other SCP studies. (D) Once the results have been submitted for publication, researchers can give reviewers anonymous, view-only access to their private SCP study. (E) When the data are ready to be shared more widely, researchers can make their SCP study public, so that anyone who visits SCP can view, explore, and download the data. (F) To further increase the impact of a line of work, researchers can collect their SCP studies into a searchable “study collection.” The cycle restarts as researchers continue exploring studies on SCP.

### Finding Data

In the early stages of research, Single Cell Portal facilitates hypothesis generation by helping researchers find and explore interesting datasets. SCP’s rich data archive makes it a valuable place to find data that are relevant to many research topics. **Figure 3** illustrates the richness of this archive, which includes public datasets on 14 species, 83 diseases, 104 organs, and 160 cell types. These studies also cover a diversity of methodologies, including 38 library preparation protocols.

**Figure 3.**
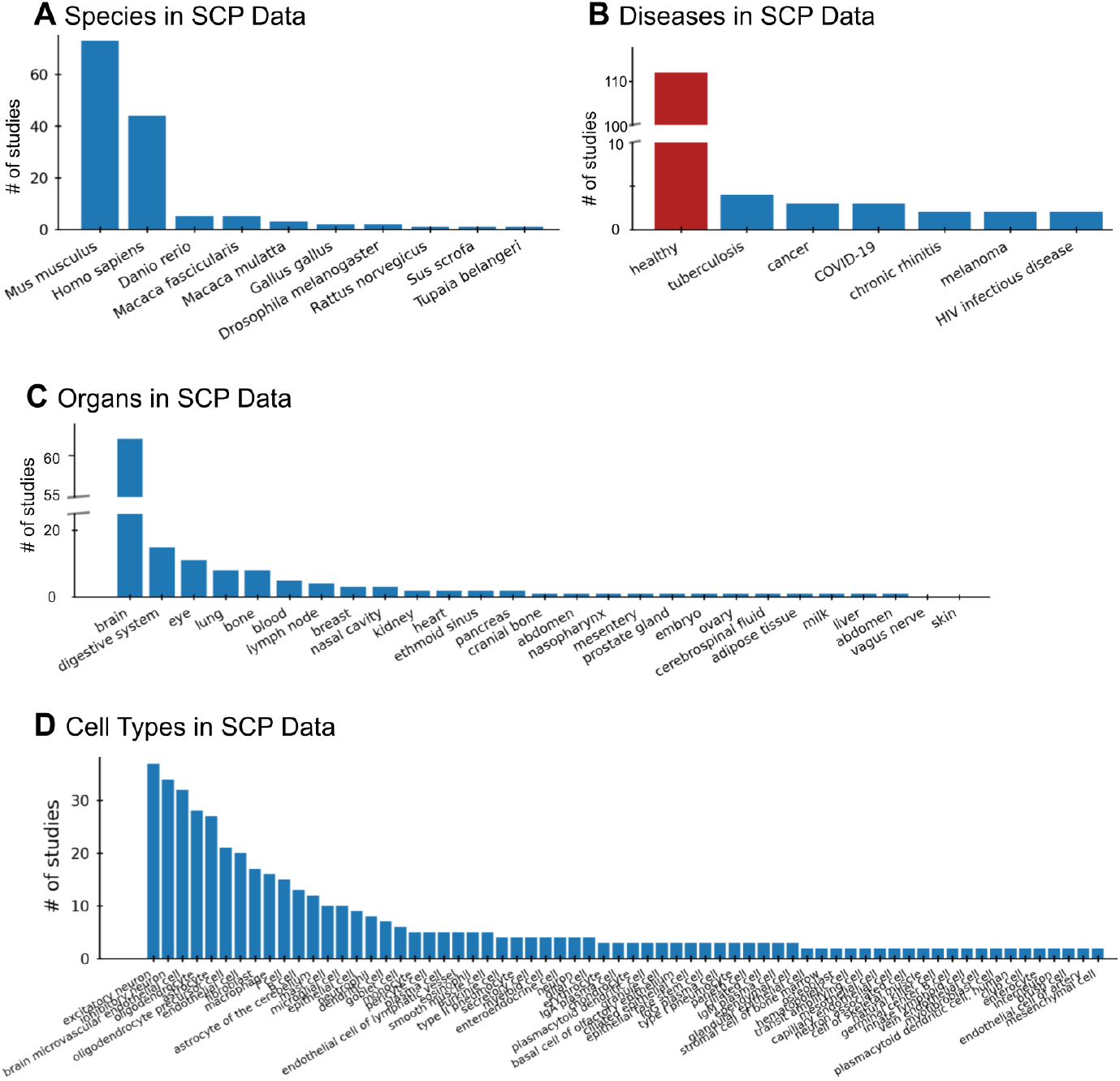
Single Cell Portal’s Rich Data Archive. Single Cell Portal attracts diverse data, making it a convenient place to investigate a variety of scientific questions. Here we plot the number of public SCP studies with data from different (A) species, (B) diseases, (C) organs, and (D) cell types. Only cell types that appeared in at least 2 studies are plotted in (D). Note that some studies include data from multiple species, diseases, or organs.

Multi-faceted search functions on SCP allow researchers to find the datasets that are most interesting to them through several means. These include searching for general information – such as a keyword, author name, or study title – and filtering for specific metadata – such as library preparation protocol, species, or cell type. In particular, SCP’s gene search feature allows researchers to search for a specific gene and then review summary plots for all studies with data on that gene. In addition to informing research hypotheses, this searchable data archive enables researchers to aggregate datasets for meta-analyses and evaluate the effectiveness of a given assay. For example, one could decide whether to use the 10x 3’ or 5’ protocol in a new experiment based on the protocol that yielded more data from a gene of interest in SCP’s public studies.

Additionally, SCP has developed cross-platform data search functionality, featuring datasets from the [Human Cell Atlas’ Data Coordination Platform](https://data.humancellatlas.org/explore/projects). This capability makes it easier for researchers to locate valuable data without separately searching through different single-cell data portals.

### Exploring Data

Once a researcher has found an interesting dataset, SCP’s suite of interactive visualization tools helps them dig further into the data. **Figure 4** provides an overview of these tools. The default plot for most SCP studies is a cell cluster plot (**Figure 4a**), which shows the overall structure of the data. Each point in these plots represents a single observation (e.g., a cell) in a low-dimensional embedding (either 2d or 3d; for example, from a UMAP projection).

**Figure 4:**
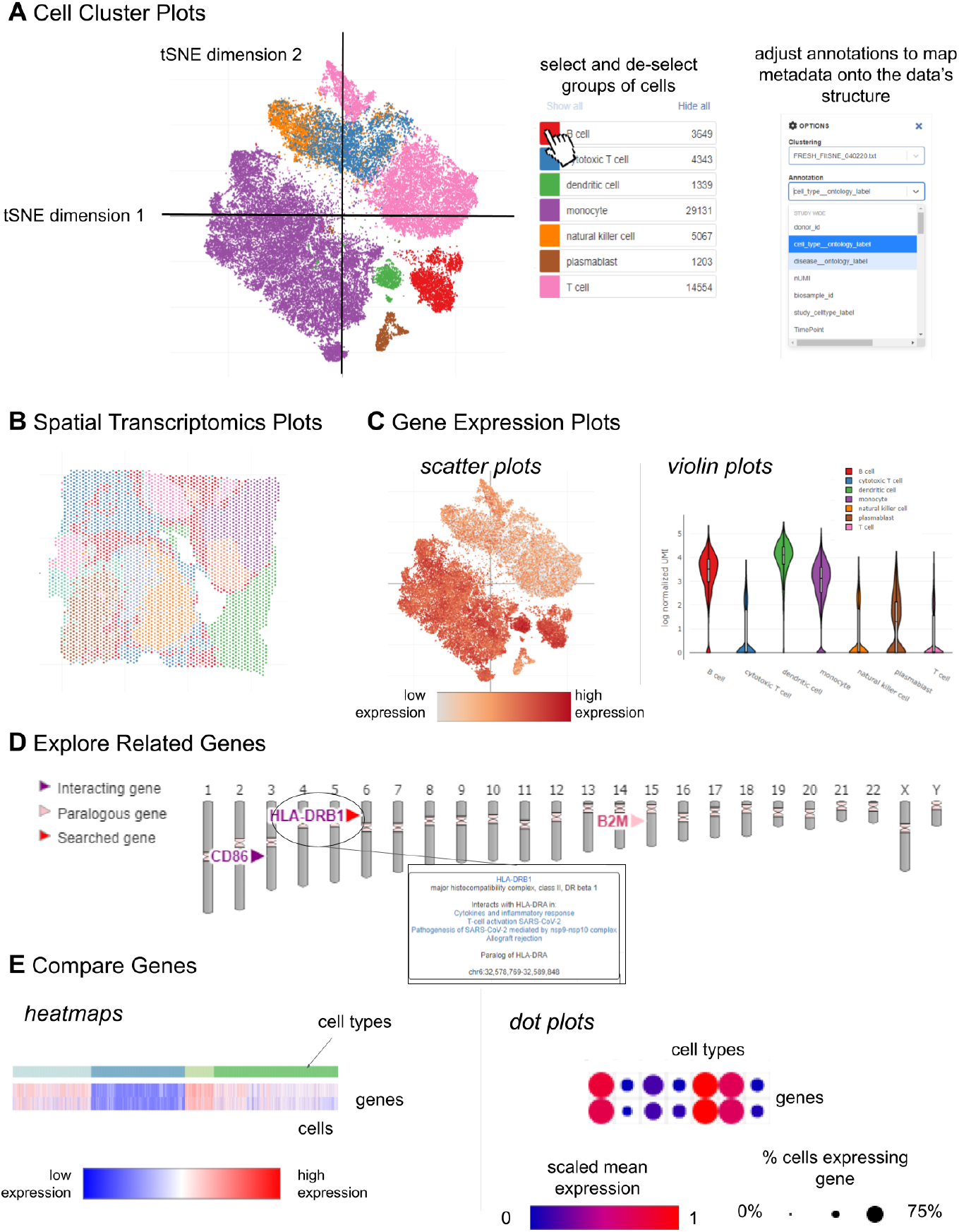
Overview of Single Cell Portal’s Interactive Visualizations. After identifying an interesting study, users can explore the data and refine hypotheses through Single Cell Portal’s interactive visualizations. (A) Cell cluster plots show the overall structure of the data and similarities between different cell types. (B) Spatial transcriptomics plots show the spatial distribution of cell types within the tissue sample. (C) Gene expression plots help users understand how expression differs across cells for a specific gene. These include gene expression scatter plots and violin plots. (D) An ideogram viewer allows users to discover genes that may be related to a gene of interest, based on the literature. (E) Heatmaps and dot plots allow users to compare genes to one another, and understand their prevalence in different cell groups.

Cells with similar genetic expression profiles are plotted close together, while cells with very different expression profiles are far apart. Each data point is colored according to an annotation, such as cell type or disease status. Users can focus on a specific subset of the data by showing or hiding cells with specific annotation labels, and can switch between several annotations and clusterings provided by the study owner.

SCP also supports plots of spatial transcriptomics data (e.g., slide-seq; [Rodrigues et al., 2019](https://www.science.org/doi/10.1126/science.aaw1219); **Figure 4b**). These plots show a spatial map of the data, and address questions about how different cell groups are distributed in the tissue. Users can view spatial plots and cell cluster plots side-by-side for a direct comparison.

To reveal the genetic mechanisms underlying the structure visible in cell cluster and spatial plots, gene expression plots show how a specific gene’s expression differs across cell types, disease states, and other variables of interest (**Figure 4c**). These include scatter plots and violin plots. Gene expression scatter plots display the same data as the cell cluster plots, but each point is colored according to the measured expression for a gene that the user has searched for, rather than an annotation such as cell type. These can be viewed side-by-side with cell cluster plots for easy comparison. Gene expression violin plots break the expression data down by cell group (e.g., cell type or disease state) to compare the overall distribution of the searched-for gene’s expression across groups.

As researchers investigate these plots to understand the contribution of a specific gene, an ideogram viewer (Weitz, Eric M. Ideogram.js. 2015) helps expand these searches to other, related genes (**Figure 4d**). After searching for a particular gene, this viewer marks interacting and paralogous genes based on findings from the literature documented in Wikipathways ([Martens et al., 2021](https://doi.org/10.1093/nar/gkaa1024)) and Ensembl ([Cunningham et al., 2022](https://academic.oup.com/nar/article/30/1/207/1332640)).

Finally, researchers can use SCP to explore differences between genes within a dataset. For most studies, this can be done by exploring heatmaps and dot plots (**Figure 4e**). In addition, a growing number of SCP’s studies include results from exploratory differential expression analyses. These are either computed by the Portal – using a nonparametric test that is most useful for identifying marker genes – or provided by the study owners, when available. These tools are particularly useful when checking patterns in researchers’ own data against other published datasets, or for spotting marker genes that are specifically expressed in one cluster of cells.

Importantly, being able to *interact* with all of these plots allows users to explore datasets more deeply than they could with a manuscript figure. In addition to the interactions described above, users can explore these plots by zooming, hovering over individual points to see more information, hovering over a legend to highlight hidden points within densely-plotted data, saving a plot as an image file, and sharing a link to a specific, user-generated plot.

### Sharing Data with Collaborators

After exploring SCP’s data and running a new experiment, researchers can deposit the cleaned, pre-processed data in a private SCP study and share the study with their collaborators (**Figure 2b**). This is a valuable step in refining the data and analyses. For example, a computational biologist ([Kedaigle, 2019](https://terra.bio/single-cell-portal-for-collaborations/)) highlighted how SCP’s user-friendly tools facilitated a recent collaboration between scientists with distinct backgrounds: “What makes SCP so valuable to me is that I can share the study workspace with my collaborators, some of whom have never installed R or heard of the command line, and they can point-and-click to explore the data without my help. Then we can discuss the results and refine the analysis together. For example, the developmental neurologists might look at the expression plots of the marker genes they know about and give me feedback on how well I’ve assigned cells to their cell types. I can then quickly update the SCP study based on their comments and send it back to them for another round of review.” ([Kedaigle, 2019](https://terra.bio/single-cell-portal-for-collaborations/)).

Such secure data sharing plays a key role in how SCP supports the single-cell research community. Computational researchers can curate a highly customized study to share with experimentalist collaborators, who can then fine-tune the data without requiring technical expertise. This interim step is available on SCP specifically because it is a self-service platform that allows users to upload data before they are finalized and publishable.

### Checking Data against Other SCP Studies

While the data are being finalized, SCP also supports the research process by providing a set of reference data for patterns that emerge in a new dataset (**Figure 2c**). For example, researchers can check their cell type annotations by looking at the dot plots showing the expression of key marker genes for other SCP studies. Similarly, any unexpected patterns in the data can be checked against comparable studies to determine whether these patterns represent a known irregularity, a glitch in the data processing, or something novel and interesting. These steps are relatively time-consuming when done through a custom analysis. Gut-checking these patterns with SCP’s visualizations may provide similar information with less effort.

### Sharing Data with Reviewers

Once a researcher has finalized their dataset and submitted a manuscript, SCP also supports the peer review process (**Figure 2d**). Researchers can grant anonymous, view-only access to their SCP study via a URL that a journal editor can pass on to reviewers without revealing their identities. The study will remain private, and reviewers will not be able to download the data or modify the study in any way. However, they will be able to explore the data through SCP’s interactive visualizations. This approach satisfies journal requirements to link a submission to a data repository, without opening the data up to public access. The researchers can then update their data based on the reviewers’ feedback before releasing it to the public.

### Publishing Data on SCP

After the review process is complete and the paper has been accepted, researchers can publish their SCP study, opening it up for anyone to explore and download (**Figure 2e**).

Publishing a study on SCP invites other researchers to engage with the data, as illustrated in **Figure 5**. Users start visiting public studies as soon as they’re released, and this interest endures for over 6 months (**Figure 5a**). SCP’s existing base of scientific users helps bolster this interest. In 2022, approximately 96,500 users visited the Portal. This number has been growing over time, as network effects attract more and more researchers to benefit from the community’s data (**Figure 5b**). Thus, a large number of users may discover a dataset on SCP, even if they haven’t read the associated paper. As users grow, so too does the number of datasets on SCP (**Figure 5c**). As of May 2023, SCP has 560 public studies, containing over 35 million cells. Sixty of these were added in 2022 alone. Study owners can track this impact for their own studies in the “usage stats” section of their study’s settings menu (**Figure 5d**).

**Figure 5:**
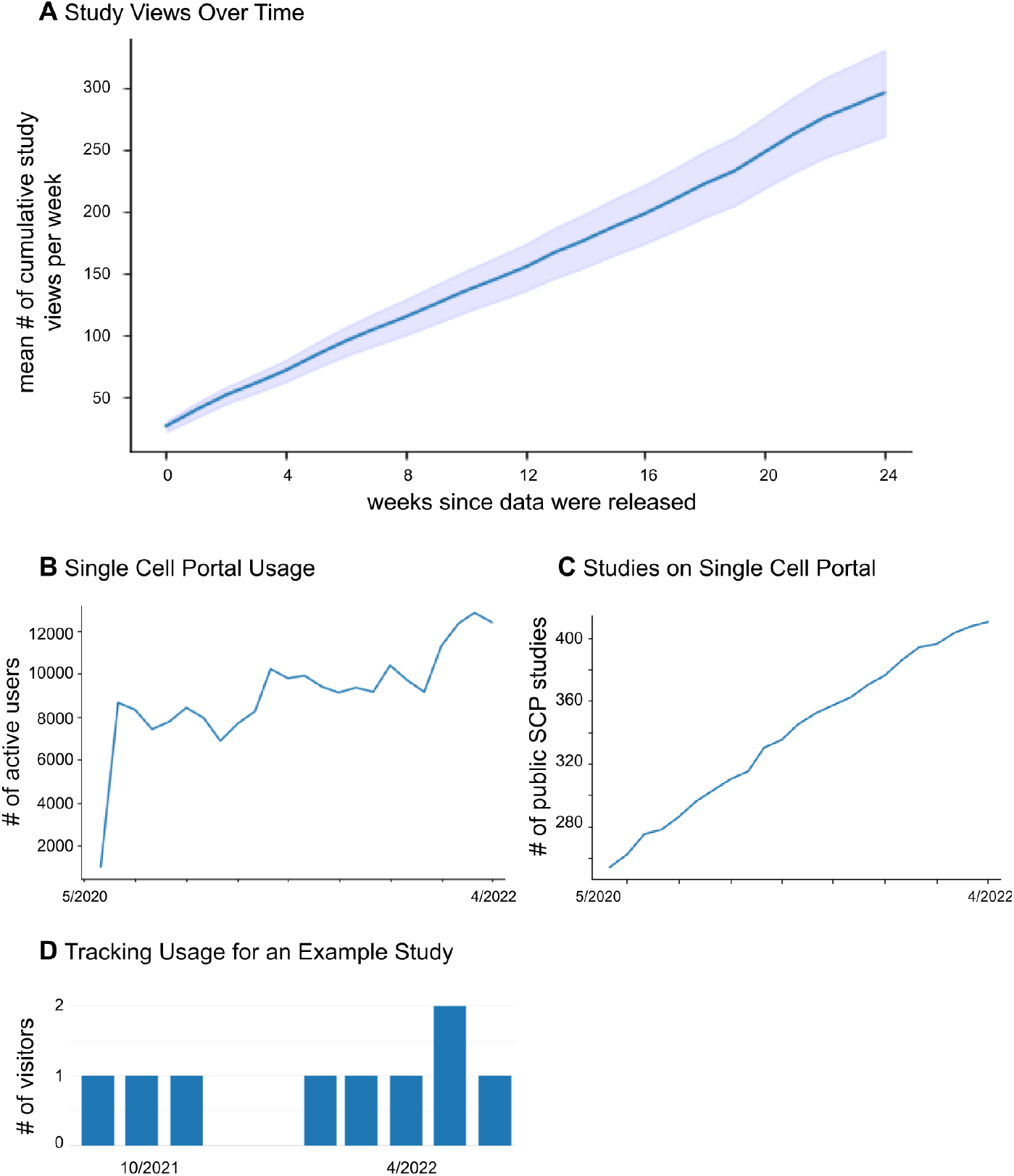
Traffic to Single Cell Portal Studies. Views of Single Cell Portal studies grow over time, as the Portal gains new users and researchers add more data. (A) Study views are plotted over time since the study went public. Cumulative study views, mean-averaged across studies, are plotted in solid blue and the standard error of this mean is shown in light-blue shading. Data are only plotted for studies that went public between August 2020 (when we began collecting study viewing data) and April 2022 (*N* = 93 studies). (B) The number of users who actively use SCP has been rising over time. User data are shown from May 2022. (C) The number of public studies available on SCP has risen steadily over time. Data for (A) and (B) were obtained from the Mixpanel analytics platform. (D) Example of plots that the study owner can use to track views and downloads for an individual study (accessible via the “study settings” menu).

To increase the impact of an SCP study and make it more discoverable, researchers can collect studies from their lab or consortium into a “study collection” (**Figure 2f**). Study collections act as a portal-within-the-Portal, showcasing studies from a particular group and allowing users to search through and explore those studies. Researchers can then link to their collection from external sites (such as a lab website) to make it easier for other researchers to explore their full body of work in one place.

## STORING DATA IN THE CLOUD

All data uploaded to SCP are stored on cloud servers as a part of the [Terra Platform](https://app.terra.bio/). Terra is a cloud-based platform for storing and analyzing biomedical data. On Terra, researchers can run their data through pipelines and notebooks that are specifically curated for their scientific quality, without having to wrangle large datasets on local machines. The results of some of these pipelines (e.g., [Cumulus](https://app.terra.bio/#workspaces/kco-tech/Cumulus); [Li et al., 2020](https://www.nature.com/articles/s41592-020-0905-x)) can be used to create an SCP study. If the results change, researchers can simply sync the changes between their SCP study and their Terra workspace.

Because of its integration with Terra, data on SCP fall within a FISMA-moderate boundary. This adds a layer of security to SCP’s data that may be required by some grants. And, to further reduce the barriers to open data sharing, most SCP data are hosted on Terra at no cost to the study owners.

## CONCLUSIONS

Single Cell Portal provides an open-access, explorable, performant way to share single-cell genomics datasets. In doing so, SCP supports researchers throughout their projects’ lifecycles. Most importantly, SCP provides interactive visualizations that make it easy for researchers to explore a large set of single-cell datasets, refine analyses with collaborators, and increase the impact of their own work on the research community. As the field of single-cell genomics continues to evolve – generating larger, more complex, and more diverse datasets – SCP will continue expanding this support to benefit as many researchers as possible.

## CODE AVAILABILITY

The source code for Single Cell Portal can be found at https://github.com/broadinstitute/single_cell_portal_core. The source code for SCP’s user-uploaded file ingest pipeline can be found at https://github.com/broadinstitute/scp-ingest-pipeline. The source code for the related genes ideogram can be found at https://github.com/eweitz/ideogram.

## ACKNOWLEDGEMENTS

Single Cell Portal has been supported by a vibrant community of scientists. Thank you to everyone who has used the portal to collaborate and communicate their science. We are grateful to Aviv Regev, Timothy Tickle, and Brian Haas for their founding vision and leadership. We thank and highlight scientists and software engineers who significantly contributed ideas and time that have shaped the portal including: Eno Akpan, Clare Bernard, Devon Bush, Jerome Chadel, Dani Chamberland, Jonah Cool, Bingxing Huo, Sarah Fortune, James Gatter, Raphael Gottardo, Imani Harrison, Naomi Habib, Matan Hofree, Vicky Horst, Velina Kozareva, Jonah Langlieb, Bo Li, Zhongjie Liu, Christine Loreth, Evan Macosko, David Mohs, Emily Munro-Ludders, Jared Nedzel, Sarah Nyquis, Yanay Rosen, Orit Rozenblatt-Rosen, Amelia Sagoff, Alex Shalek, Karthik Shekhar, Lou Strano, Charles Vanderburg, Kyle Vernest, Kendra West, and Nir Yosef.

Funding for Single Cell Portal has been generously provided by the Klarman Family Foundation as a part of the Klarman Cell Observatory, the Bill and Melinda Gates Foundation grant #OPP1202327 and grant #INV-027498, NIH Grant U19MH114821-05, and NIH Grant R24MH114788.

## Notes

### Competing Interest Statement

The authors have declared no competing interest.

